# Deuterated buprenorphine retains pharmacodynamic properties of buprenorphine and resists metabolism to the active metabolite norbuprenorphine in rats

**DOI:** 10.1101/2022.12.12.520120

**Authors:** Venumadhav Janganati, Paloma Salazar, Brian J. Parks, Gregory S. Gorman, Paul L. Prather, Eric C. Peterson, Alexander W. Alund, Jeffery H. Moran, Peter A. Crooks, Lisa K. Brents

## Abstract

An active metabolite of buprenorphine (**BUP**), called norbuprenorphine (**NorBUP**), is implicated in neonatal opioid withdrawal syndrome when BUP is taken during pregnancy. Therefore, reducing or eliminating metabolism of BUP to NorBUP is a novel strategy that will likely lower total fetal exposure to opioids and thus improve offspring outcomes. Precision deuteration alters pharmacokinetics of drugs without altering pharmacodynamics. Here, we report the synthesis and testing of deuterated buprenorphine (**BUP-D2**). We determined opioid receptor affinities of BUP-D2 relative to BUP with radioligand competition receptor binding assays, and the potency and efficacy of BUP-D2 relative to BUP to activate G-proteins via opioid receptors with [^35^S]GTPγS binding assays in homogenates containing the human mu, delta, or kappa opioid receptors. The antinociceptive effects of BUP-D2 and BUP were compared using the warm-water tail withdrawal assay in rats. Blood concentration versus time profiles of BUP, BUP-D2, and NorBUP were measured in rats following intravenous BUP-D2 or BUP injection. The synthesis provided a 48% yield and the product was ≥99% deuterated. Like BUP, BUP-D2 had sub-nanomolar affinity for opioid receptors. BUP-D2 also activated opioid receptors and induced antinociception with equal potency and efficacy as BUP. The maximum concentration and the area under the curve of NorBUP in the blood of rats that received BUP-D2 were over 19- and 10-fold lower, respectively, than in rats that received BUP. These results indicate that BUP-D2 retains key pharmacodynamic properties of BUP and resists metabolism to NorBUP and therefore holds promise as an alternative to BUP.

**GRAPHICAL ABSTRACT:** 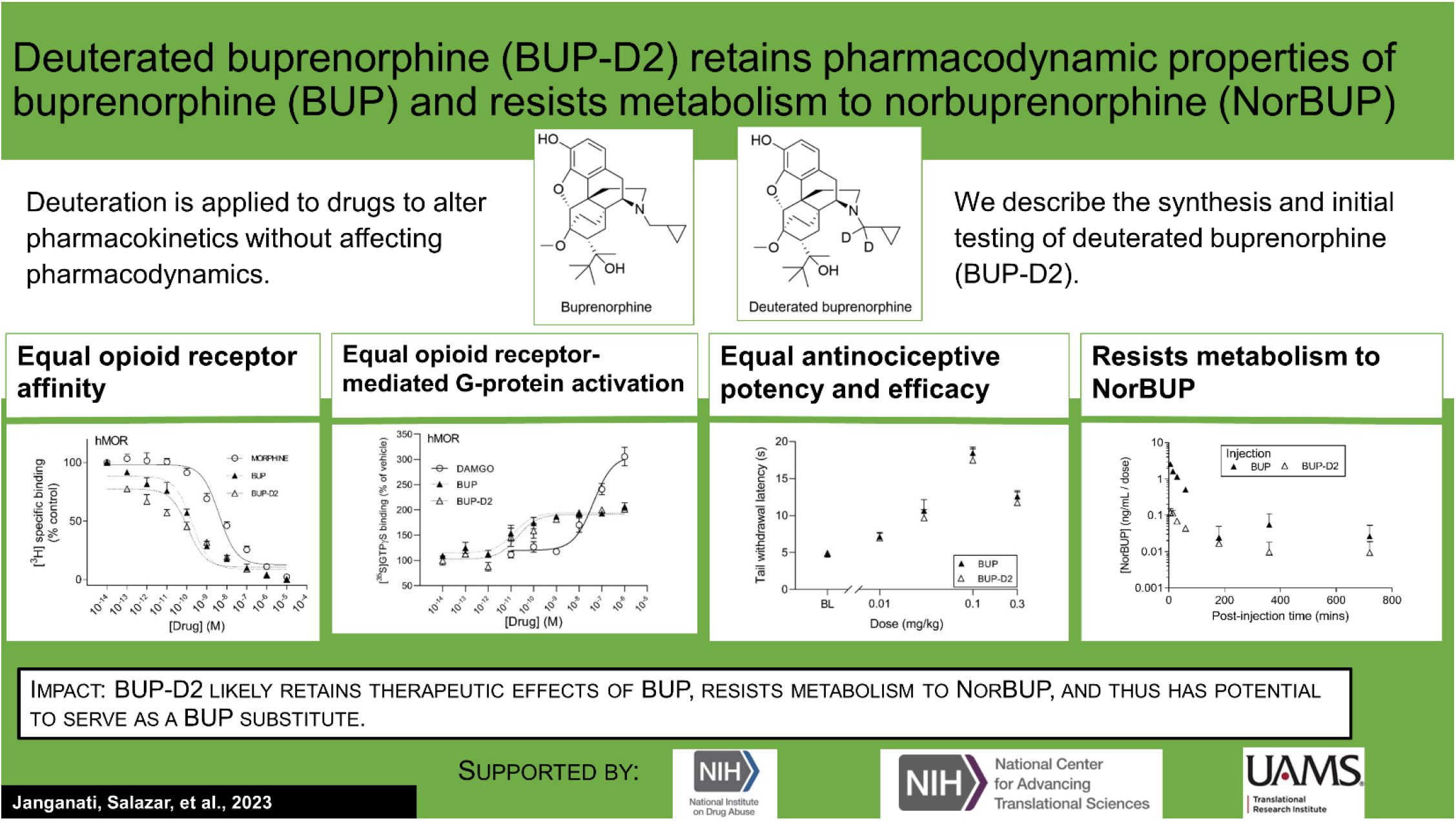

## 1. INTRODUCTION

Chronic use of opioids during pregnancy negatively affects mothers and their children, and prenatal opioid use has increased more than five-fold in the United States since the beginning of the opioid addiction and overdose crisis^1,2^. Neonatal opioid withdrawal syndrome (**NOWS**) is a major adverse consequence of prenatal opioid exposure. NOWS is defined by multisystem dysregulation, resulting in poor feeding, vomiting, diarrhea, sweating, lacrimation, tremors, seizures, hyperalgesia, poor sleep, extreme irritability, and sensitivity to lights and sounds^3^. This complex syndrome interferes with growth and development of the neonate and often requires care in costly neonatal intensive care units for the first days, and even weeks, of life^4^. In addition to NOWS, prenatal opioid use is associated with other adverse birth outcomes, including maternal death^5^, stillbirth^5,6^, preterm labor^5^/birth^6–8^, premature rupture of membranes^5^, intrauterine growth restriction^5,7^, placental abruption^5^, low birth weight (< 2500 g)^8^, being small for gestational age (weight < 2 standard deviations below sex- and gestational age-specific mean)^8^, and having congenital malformations^8^.

Maternal opioid use disorder (**OUD**) drives chronic opioid use, high stress, and poor self-care during pregnancy, but can be treated with the mu opioid receptor agonists methadone or buprenorphine (**BUP**) as part of medication-assisted treatment. Medication-assisted treatment stabilizes the maternal-fetal dyad by preventing opioid withdrawal and craving, thus diminishing maternal motivation to pursue and use potentially dangerous illicit opioids. Duration of prenatal medication-assisted treatment is associated with decreased risk of overdose^9^, preterm birth^9^, and low birth weight^9^. However, because methadone and BUP are opioids, medication-assisted treatment with them during pregnancy is associated with NOWS^9^. Prenatal treatment with BUP yields better neonatal outcomes than prenatal treatment with methadone, including shorter hospital stays and less medication needed to treat withdrawal^10^. However, because BUP is a partial agonist of mu opioid receptors (**MOR**), the receptor subtype that mediates most clinically sought effects of opioids, chronic use of BUP often causes dependence and leads to withdrawal upon cessation of exposure. As such, approximately half of children born to women treated with BUP develop moderate-to-severe NOWS that requires weaning with an opioid agonist such as morphine or methadone^10,11^. Although prenatal BUP treatment currently produces the best outcomes for neonates of women with OUD, improved treatments are clearly needed.

Multiple lines of evidence suggest that an active metabolite of BUP called norbuprenorphine (**NorBUP**) contributes to the adverse effects of BUP on the fetus. NorBUP is a major metabolite of BUP that is formed by *N*-dealkylation by multiple cytochrome 450 enzymes (CYPs), predominantly CYPs 3A4, 3A5, 2C8, and 19A1^12–14^. In contrast to the partial agonist activity of BUP, NorBUP is a high efficacy mu opioid receptor agonist^15^; thus, NorBUP activates mu opioid receptors with efficacy that more closely resembles that of methadone and other high efficacy opioids that are associated with more severe NOWS than BUP. Following chronic prenatal BUP use, concentrations of NorBUP in the human placenta, umbilical cord plasma, and meconium are at least 10-fold higher than BUP concentrations^16–18^, suggesting that fetal exposure to NorBUP exceeds BUP. NOWS severity is independent of maternal BUP dose^16,19,20^, suggesting that factors other than dosing (*e.g*., individual variability in BUP metabolism) determine fetal exposure to BUP and its active metabolites. For example, it is plausible that NOWS is more severe for neonates exposed *in utero* to higher levels of NorBUP due to dominance of the NorBUP metabolic pathway over alternative pathways, such as glucuronide conjugation. This possibility is supported by the results of a study in which investigators measured umbilical cord blood concentrations of BUP and its metabolites (which represents fetal blood concentrations at delivery) and determined that NorBUP, but not BUP, was positively correlated with NOWS severity^16^. Furthermore, the glucuronide conjugate of BUP was negatively correlated with NOWS severity^16^, suggesting that bias for glucuronide conjugation (over *N*-dealkylation to form NorBUP) is protective, and bias for NorBUP formation (over glucuronide conjugation) is more harmful. Additionally, we previously showed with a rat model of NOWS that prenatal exposure to NorBUP induces dependence and neonatal withdrawal at levels comparable to morphine^21^. NorBUP likely contributes little to the therapeutic centrally mediated effects of BUP treatment due to its restriction to the periphery by p-glycoprotein in the blood-brain barrier^22,23^. Altogether, this evidence suggests that decreasing fetal exposure to NorBUP by reducing NorBUP formation can potentially improve short-term neonatal outcomes, such as NOWS, and the long-term effects of opioid exposure on neurodevelopment.

In this study, we applied precision deuteration to BUP and determined whether deuteration affected its pharmacodynamic properties, which included opioid receptor affinity and potency and efficacy to activate G-proteins via opioid receptors. We also investigated its pharmacokinetic properties with specific interest in how deuteration affected NorBUP blood concentrations.

Precision deuteration involves replacing hydrogen atoms with their heavy isotope deuterium, which exploits the primary kinetic isotope effect to slow bond-breaking by metabolic enzymes^24^ (**Fig. 1**). The expected net result of this small change is altered pharmacokinetics (*e.g*., slowing and reduction of metabolite formation) with little or no change in pharmacodynamics^25,26^. In the present work, we synthesized a deuterated buprenorphine (**BUP-D2**) by replacing two methylene hydrogens with deuterium atoms adjacent to the tertiary nitrogen in the buprenorphine structure (**Fig. 2**) to inhibit oxidative dealkylation of the *N*-cyclopropylmethyl moiety by liver cytochrome P450’s. We hypothesized that BUP and BUP-D2 have equal affinity, potency, and efficacy for opioid receptors, have equal potency and efficacy of antinociception, and that blood concentrations of NorBUP are lower in rats following injection with BUP-D2 relative to BUP. Here, we report the synthesis and initial testing of BUP-D2, compared to BUP, for opioid receptor affinity and G-protein activation using a radioligand competition receptor binding and [^35^S]GTPγS binding assays, respectively. We then tested BUP-D2 and BUP in the warm water tail withdrawal assay, a rat model of antinociception, to determine whether precision deuteration alters *in vivo* buprenorphine effects. Last, we compared NorBUP blood concentration-time profiles up to 12 hours after bolus intravenous administration of BUP-D2 relative to BUP of rats. We determined that BUP and BUP-D2 have virtually indistinguishable pharmacodynamic properties and that NorBUP blood concentrations were over 10-fold lower in BUP-D2-treated rats versus BUP-treated rats. The results of the present study suggest that BUP-D2 shows promise as an alternative to BUP for treating OUD.

**Figure 1.**
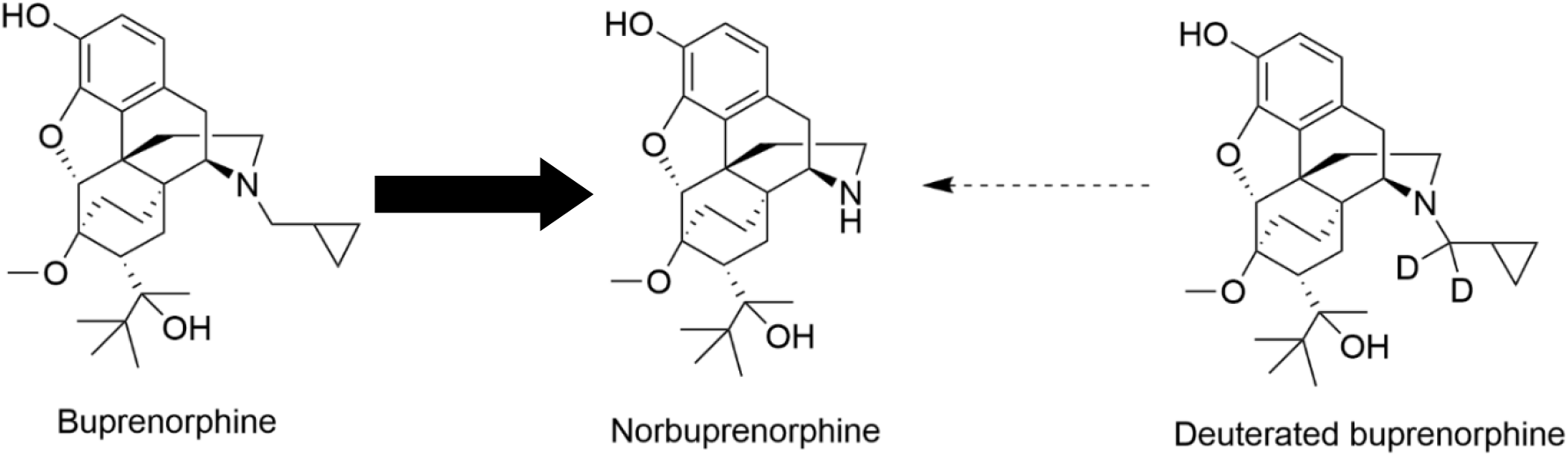
Norbuprenorphine formation. In humans, norbuprenorphine is the primary major metabolite of buprenorphine, as represented by the bold arrow. Norbuprenorphine is formed by oxidative *N*-dealkylation of buprenorphine’s cyclopropylmethyl group by various cytochrome P450s. Precision deuteration is expected to make this group less vulnerable to metabolic cleavage and thus reduce its metabolism to norbuprenorphine, as represented by the thin broken arrow.

**Figure 2.**
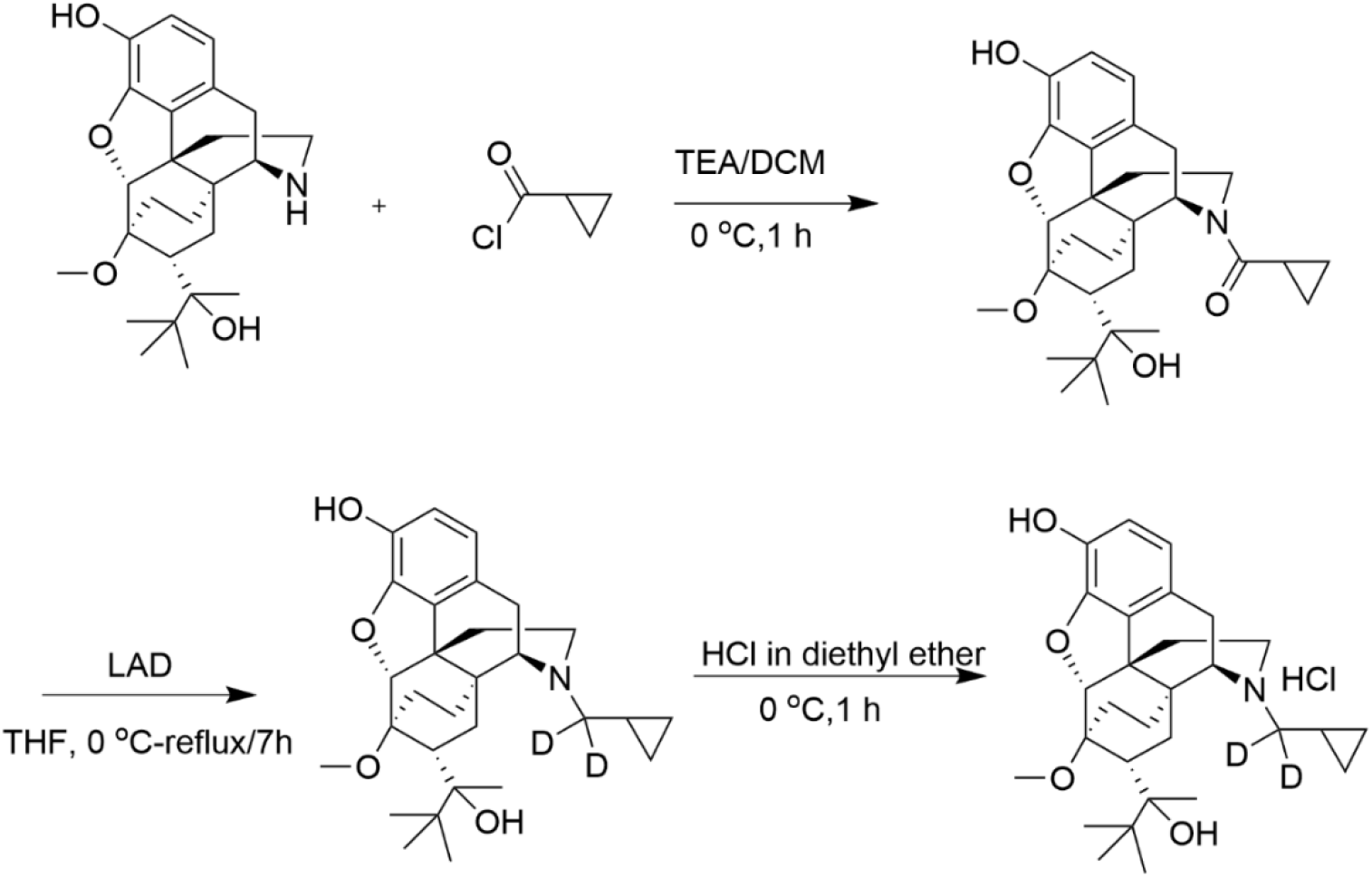
Synthesis of BUP-D2 Hydrochloride. See Materials and Methods for description of synthesis.

## 2. MATERIALS AND METHODS

### 2.1 Drugs

The National Institute on Drug Abuse Drug Supply Program (Rockville, MD) provided norbuprenorphine (**NorBUP**) in free base form. Buprenorphine (**BUP**) was purchased from Tocris (Minneapolis, MN). Morphine sulfate salt pentahydrate was purchased from Sigma-Aldrich (St. Louis, MO). With the exception of NorBUP, solvents and chemicals used to synthesize deuterated buprenorphine were purchased from Sigma-Aldrich (St. Louis, MO). [^3^H]diprenorphine (25.1 Ci/mmol) and [^35^S]GTPγS (1250 Ci/mmol) were purchased from Perkin-Elmer (Waltham, MA). Guanosine-5-diphosphate (GDP) and unlabeled GTPγS were purchased from Fisher Scientific (Waltham, MA) and Sigma-Aldrich (St. Louis, MO), respectively. [D-Ala^2^, NMe-Phe^4^, Gly-ol^5^]-enkephalin (**DAMGO**) was purchased from Cayman Chemical (Ann Arbor, MI). [D-Pen^2,5^]Enkephalin, [D-Pen^2^,D-Pen^5^]Enkephalin (**DPDPE**) and racemic U-50,488 HCl were purchased from Tocris (Minneapolis, MN). Naltrexone hydrochloride was purchased from Sigma Aldrich (St. Louis, MO). Drugs for *in vitro* assays were dissolved at concentrations of 10^-2^ M in 100% DMSO and stored at −20°C until used in experiments. Drugs for *in vivo* experiments were dissolved in a sterile saline solution (0.9% NaCl) containing 5% DMSO. ToxBox^®^ analytical plates (PinPoint Testing, LLC, Little Rock, AR) were used to quantify BUP, BUP-D2, and NorBUP in rat blood (see Section 2.9 for details).

### 2.2 BUP-D2 synthesis

BUP-D2 was synthesized in a two-step process as illustrated in **Figure 2** and described here:

**Step 1.** *Cyclopropyl((4R,4aS, 6R, 7R, 7aR, 12bS)-9-hydroxy-6-(2-hydroxy-3,3-dimethylbutan-2-yl)-7-methoxy-1,2,5,6,7,7a-hexahydro-4a, 7-ethano-4,12-methanobenzofuro[3,2-e]isoquinolin-3(4H)-yl)methanone:* Triethylamine (44.96 mg, 0.444 mmol) and cyclopropanecarbonyl chloride (23.22 mg, 0.222 mmol) were added to norbuprenorphine hydrochloride (100 mg, 0.222 mmol) dissolved in dichloromethane (3 mL) at 0 °C. The reaction mixture was stirred for 1 h at 0 °C. After completion of the reaction, which was monitored by thin layer chromatography (**TLC**), water (5 mL) was added, and the reaction mixture was extracted with dichloromethane (2×5 mL). The organic layer was washed with water (3×5 mL), dried over anhydrous Na2SO4 and concentrated under reduced pressure to afford crude product. The crude product was purified by column chromatography (silica gel, 30-40 % EtOAc in hexane) and obtained as a mixture of diastereomers (yield: 75 %).
**Step 2.** (*4R,4aS,6R, 7R, 7aR, 12bS)-3-(cyclopropylmethyl-d_2_)-6-(2-hydroxy-3,3-dimethylbutan-2-yl)-7-methoxy-1,2,3,4,5,6,7,7a-octahydro-4a, 7-ethano-4,12-methanobenzofuro[3,2-e]isoquinolin-9-ol:* To the cyclopropylcarbonyl norbuprenorphine (diasteriomeric mixture) (80 mg, 0.17 mmol) in tetrahydrofuran (3 mL), lithium aluminum deuteride (27.92 mg, 0.68 mmol) was added at 0 °C. The reaction mixture was stirred at 50 °C for 3 h. After completion of the reaction, which was monitored by TLC, the reaction mixture was brought to 0 °C and quenched by slow addition of 0.03 mL of water, followed by 0.03 mL of 15% aqueous sodium hydroxide, and 0.03 mL of water; the resulting mixture was then warmed to ambient temperature, stirred for 30 min and filtered. The reaction mixture was then stirred for 30 min at ambient temperature and filtered. The filtrate was concentrated under reduced pressure to afford crude product. The crude product was further purified by flash column chromatography (silica gel, 30 % EtOAc in hexane) to afford the pure product as a white solid. The solid was dissolved in diethyl ether and cooled to 0 °C. 2M HCl solution in diethyl ether (~ 0.1 mL) was added and the resulting mixture stirred for 1 h. The reaction mixture was filtered and washed with diethyl ether to afford a white solid (yield: 64 %).

^1^H and ^13^C NMR spectra were recorded on a Varian 400 MHz spectrometer equipped with a Linux workstation running on vNMRj software. All the spectra were phased; baseline was corrected where necessary, and solvent signals (CDCl3) were used as reference for both ^1^H and ^13^C spectra. TLC separations were carried out on pre-coated silica gel plates (F 254 Merck).

^1^H NMR (400 MHz, CDCl_3_; **Supporting Data Figure 1A**): *δ* 6.81 (d, *J* = 7.6 Hz, 1H), 6.50 (d, *J* = 7.6 Hz, 1H), 5.81 (s, 1H), 4.51 (s, 1H), 4.44 (s, 1H), 3.51 (s, 3H), 2.98-2.83 (m, 3H), 2.62-2.55 (m, 1H), 2.32-2.12 (m, 3H), 2.01-1.92 (m, 1H), 1.85-1.64 (m, 3H), 1.54 (s, 1H), 1.34 (s, 3H), 1.31-1.25 (m, 1H), 1.1-1.0 (m, 9H), 0.81-0.62 (m, 2H), 0.51-0.41 (m, 2H), 0.12-0.06 (m, 2H) *ppm*. ^13^C NMR (100 MHz, CDCl_3_; **Supporting Data Figure 1B**): *δ* 145.4, 137.3, 132.5, 128.2, 119.5, 116.4, 96.9, 80.8, 79.6, 58.2, 52.5, 46.4, 43.6, 40.3, 35.9, 35.6, 33.4, 29.6, 26.3, 22.9, 20.1, 18.1, 9.2, 4.0, 3.1 *ppm*. MS (ESI), m/z: 470 (M+H)^+^.

Enrichment of the product with deuterium was determined by LC/MS/MS to be >99% g/atom (**Supporting Data Figure 2**). The LC/MS/MS system used for this analysis consisted of a Shimadzu system (Columbia MD) equipped with LC20-AD dual HLPC pumps, a SIL20-AC HT autosampler, and a DGU-20A2 in-line degasser. Detection was performed using an Applied BioSystems 4000 QTRAP (Applied Biosystems, Foster City, CA) triple quadrupole mass spectrometer operated in the positive ion mode. Mass calibration, data acquisition and quantitation were performed using Applied Biosystem Analyst 1.6.21 software (Applied Biosystems, Foster City, CA).

### 2.3 Cell Culture

Chinese hamster ovary (**CHO**) cells stably transfected with human mu opioid receptors (**CHO-hMOR**) were a gift from Dr. Dana Selley (Virginia Commonwealth University, Richmond, VA). CHO cells were stably transfected with human delta opioid receptors (**CHO-hDOR**) in our laboratory as described previously^27^. CHO cells stably transfected with human kappa opioid receptors (**CHO-hKOR**) were a gift from Dr. Lee-Yuan Liu-Chen (Temple University, Philadelphia, PA). CHO-hMOR and CHO-hDOR cells were cultured in DMEM Nutrient Mix F12 1:1 media containing 10% FetalPlex^®^, 1% penicillin-streptomycin, and 0.2 mg/mL hygromycin (Gibco, Waltham, MA). CHO-hKOR cells were cultured in DMEM Nutrient Mix F12 1:1 (Gibco, Waltham, MA) media containing 10% FetalPlex^®^ (Gemini Bioproducts, Sacramento, CA), 1% penicillin-streptomycin (Corning, Corning, NY), and 0.27 mg/mL G418 Sulfate (Fisher Scientific, Houston, TX). Cells were grown in a humidified chamber with 5% CO_2_ at 37°C in sterile T175 flasks and were harvested at 100% confluence with 0.04% EDTA in phosphate buffered saline (pH 7.5). Harvested cells were centrifuged at 1040 *x g* for 10 mins to form cell pellets and supernatant was removed. Cell pellets were stored at −80°C until used to make membrane homogenates.

### 2.4 Membrane Homogenate Preparation

Membrane homogenates were prepared as follows for the experiments described in Section 2.5 and 2.6. Up to five cell pellets, each made from up to four 100% confluent T175s, were thawed on ice, then transferred to a 40 mL Dounce glass homogenizer and suspended in 10 mL (or ~2 mL/cell pellet) of ice-cold homogenization buffer (50 mM HEPES, pH 7.4, 3 mM MgCl_2_, and 1 mM EGTA). Ten strokes were applied to the contents of the homogenizer with a coarse-grade pestle, and the contents were then centrifuged at 40,000 *x g* for 10 minutes at +4°C. Supernatants were discarded, and the pellet was again transferred to the homogenizer, suspended in 10 mL of homogenization buffer, coarsely homogenized with 10 strokes, and centrifuged at 40,000 *x g* for 10 minutes at +4°C twice more, with the supernatant discarded each time. The resulting pellet was homogenized using 10 strokes of a fine-grade pestle in 10 mL (or ~2 mL/cell pellet) of ice-cold 50 mM HEPES. Homogenates were aliquoted in 0.5 or 1 mL volumes and stored at −80°C until used in experiments. Protein concentrations of homogenates were determined using the BCA Protein Assay (Thermo Scientific, Walham, MA).

### 2.5 Competition Receptor Binding

Competition receptor binding assays were performed using 25, 50, or 100 μg of membrane homogenates (prepared as described in Section 2.4) made from CHO-hMOR, CHO-hKOR, or CHO-hDOR cells, respectively, per sample. Membranes were incubated with 1 nM of the non-selective opioid antagonist [^3^H]diprenorphine plus vehicle or naltrexone (10 μM, non-specific binding) or various concentrations of either non-radioactive morphine, buprenorphine, or deuterated buprenorphine in a buffer containing 5 mM MgCl_2_, 50 mM Tris-HCl, and 0.05% bovine serum albumin. Total reaction volume was 1 mL. Samples were incubated for 90 minutes at room temperature to achieve equilibrium before reactions were rapidly terminated by vacuum filtration onto Whatman GF/B glass microfiber filters (Brandel, Gaithersburg, MD) with a Brandel cell harvester (Brandel, Gaithersburg, MD). Filters were washed three times with 5 mL of ice-cold assay buffer and samples were transferred to 7-mL scintillation vials. Four milliliters of ScintiVerse™ BD Cocktail scintillation fluid (Fisher Scientific, Waltham, MA) were added to scintillation vials, and allowed to incubate overnight. Radioactivity retained on filters was then quantified by liquid scintillation spectrophotometry using a Perkin-Elmer Tri-Carb 2910TR Liquid Scintillation Analyzer (Walham, MA). Non-specific binding was determined in each experiment using a receptor-saturating concentration of non-radioactive naltrexone, a non-selective opioid antagonist. Specific binding for each sample was determined by subtracting radioactive counts (decays per minute, **DPMs**) measured in these non-specific binding samples from the total radioactive counts obtained in samples. Samples were conducted in duplicate within each experiment, and at least three independent experiments were combined to produce each receptor binding curve.

### 2.6 [^35^S]GTPγS binding assay

Activation of MOR, DOR, and KOR by BUP and BUP-D2 was quantified using an [^35^S]GTPγS, a radiolabeled non-hydrolyzable analogue of GTP that binds and labels activated G-proteins. In this assay, 50 μg of membrane homogenates (prepared as described in Section 2.4) were incubated with 0.1 nM [^35^S]GTPγS in the presence of vehicle, unlabeled GTPγS (10 μM, non-specific binding) or opioids (BUP, BUP-D2, or controls) for 30 minutes at 30°C. The receptor-selective full agonists DAMGO, U50, 488, and DPDPE were used as positive controls to measure activation of hMOR, hKOR, and hDOR, respectively. Naltrexone, a non-selective opioid antagonist, was included as a negative control. Assay buffer contained 20 mM HEPES, 100 mM NaCl, 10 mM MgCl_2_, 10 μM GDP, and 0.05% bovine serum albumin. Membrane homogenates and GDP were pre-incubated together for 5 minutes at room temperature before being combined with the other components. Reactions were rapidly terminated using vacuum filtration onto Whatman GF/B glass microfiber filters (Brandel, Gaithersburg, MD) using a Brandel cell harvester, followed by three 5 mL washes of ice-cold 50 mM Tris-HCl (pH 7.4) buffer containing 0.05% bovine serum albumin. Filters were transferred to 7 mL scintillation vials and 4 mL of ScintiVerse™ BD Cocktail scintillation fluid was added to each vial. Following overnight incubation, bound radioactivity in samples was determined by liquid scintillation spectrophotometry using a Perkin-Elmer Tri-Carb 4910 TR Liquid Scintillation Analyzer (Walham, MA). Non-specific binding was determined in each experiment using a receptor-saturating concentration of non-radioactive GTPγS. Specific binding for each sample was determined by subtracting radioactive counts (DPMs) measured in these non-specific binding samples from the radioactive counts in the sample. Samples were conducted in triplicate within each experiment, and at least three independent experiments were combined to produce each G-protein activation curve.

### 2.7 Animal Care and Use

Studies with rodents were approved by the University of Arkansas for Medical Sciences Institutional Animal Care and Use Committee (IACUC) prior to commencement of experiments. Female Long Evans rats with indwelling jugular catheters were purchased from Charles River Laboratories (Wilmington, MA). Female rats weighed 200-250 grams and were 7-10 weeks old upon arrival. All rats were singly housed in Plexiglass cages with enrichment. Rats were kept on 14-hour light/10-hour dark cycles with free access to food in a temperature and humidity-controlled room.

### 2.8 Warm-water tail withdrawal assay

The analgesic activity of BUP and BUP-D2 were compared using the warm-water tail withdrawal procedure in rats during the light phase of the light-dark cycle (1200-1600). A total of 29 rats were used in this experiment.

In this procedure, the latency of a rat to remove its tail from warm water (50°C) up to a cutoff of 20 seconds was measured as an endpoint of nociception. Control trials using a non-noxious water temperature (40°C) were conducted between test trials to confirm that tail withdrawals during test trials were specifically a nociceptive response (as opposed to a learned response). Prior to testing, animals were acclimated to the procedure with control trials until they had at least two successful control trials (*i.e*., trials in which animals did not remove tail from non-noxious water before cutoff). Then a test trial was conducted using 50°C water to determine baseline latency to withdraw the tail. Ten minutes later rats were given an intravenous injection via jugular catheter of either BUP or BUP-D2 (0.01, 0.03, 0.1, or 0.3 mg/kg, 2 ml/kg). Additional test trials were conducted 10 and 60 minutes after dosing. Control trials were conducted 5 minutes before dosing and 5 and 30 minutes after dosing. The investigator conducting the test was blinded to treatments.

### 2.8 Pharmacokinetic Experiment

A total of six rats were used in this experiment. Rats were lightly anesthetized with isoflurane carried by 100% O2 (3-4% at 1.5 L/min) before administration of 2 mL/kg bolus tail vein injections containing either BUP HCl (0.9 mg/kg) or BUP-D2 HCl (5.6 mg/kg) in 5% DMSO dissolved in sterile saline solution (0.9% NaCl). Blood samples (250 μL) were taken from each rat via jugular catheter at the following time points after drug administration: 6, 15, 30, 60, 180, 360, and 720 mins. Samples were immediately transferred to BD Microtainer tubes containing EDTA, inverted 10 times, and stored at 4°C until LC/MS/MS analysis.

### 2.9 LC/MS/MS Analysis

Concentrations of NorBUP, BUP, and BUP-D2 were quantified in rat blood using supported liquid extraction and customized, commercially available ToxBox^®^ analytical designed by PinPoint Testing, LLC, similar to our previous work described in Griffin et al, 2019. Calibration standards and second-source quality control material were prepared on the ToxBox^®^ analytical plates by adding blank rat blood samples (250 μL whole blood) to wells containing pre-titrated NorBUP, BUP, and BUP-D2 ranging from 1.25 to 250 ng/mL and NorBUP-D3 (internal standard for NorBUP) and BUP-D4 (internal standard for BUP and BUP-D2). Experimental samples (250 μL) were added to ToxBox^®^ analytical plate wells containing only internal standards and were processed identical to standards and quality controls. Plates were shaken at room temperature for 15 mins at 900 RPMs, followed by addition of 250 μL of 0.5 M ammonium hydroxide and 15 mins of additional shaking at 900 RPMs at room temperature. Samples were then loaded onto a 96-well ISOLUTE SLE+ plate (Biotage, Charlottle, NC) and extracted using two elutions of ethyl acetate (100%, 2 × 900 μL) under gentle vacuum. The eluent was dried under nitrogen flow with gentle heating and reconstituted in 100 μL of 100% methanol. Analysis was completed using an Agilent 1260 series quaternary liquid chromatograph system that was interfaced with Agilent 6420A tandem mass spectrometer (Agilent Technologies, Santa Clara, CA). The LC-MS-MS method used a Kinetex 2.6 μm Phenyl-Hexyl 100Å LC Column (50 x 4.6 mm; Phenomenex, Inc., Torrance, CA) heated to 35°C. Analytes and internal standards were resolved using a 10 mM ammonium formate/0.1% formic acid in methanol gradient started at 95% aqueous, ramped up to 100% organic over 4 mins and held constant for 1 additional min. The gradient was then returned to initial conditions and equilibrated for an additional 2 mins. The total run time for each sample analysis was 7 min, including the column equilibration period between injections.

### 2.10 Statistical analysis

Analyses were performed using GraphPad Prism (Version 8.0; San Diego, CA). After baseline and/or non-specific values were subtracted and normalization corrections were applied, data were fit using “three-parameter agonist (or inhibitor) vs. response” non-linear regression (for competition receptor binding and [^35^S]GTPγS binding assays) or linear regression (for the warm-water tail withdrawal assay). In the competition receptor binding experiments, the log-transformed IC50 value of each replicate was obtained from the non-linear regression analyses and converted to a log-transformed K_i_ value (a normally distributed quantitative measure of receptor affinity) by applying the Cheng-Prusoff equation^28^. Statistically significant differences in affinity for each receptor subtype among BUP, BUP-D2, and morphine were determined by one-way ANOVA and Tukey’s Multiple Comparison *post-hoc* test (p < .05). In the concentration-effect [^35^S]GTPγS binding experiments, log-transformed EC50 values and log-transformed 95% confidence intervals were derived from non-linear regression analyses applied to the aggregate data. Lack of overlap of log-transformed confidence intervals, within receptor subtype, indicated statistically significant group differences in receptor potency. “Top” values (and their 95% confidence intervals) calculated by GraphPad Prism from the agonist vs. response non-linear regression analyses are reported here as the maximum effect (E_max_) values. For single-concentration [^35^S]GTPγS binding experiments, data are reported as percent vehicle control, and a one-way ANOVA with Tukey’s multiple comparisons test was conducted within receptor subtype. The ED_50_ of antinociception in the warm-water tail withdrawal assay was determined by first converting tail withdrawal latencies to percent maximum possible effect (**% MPE**) using the following formula: % MPE = (X – baseline) / cutoff – baseline) * 100, wherein “X” equals tail withdrawal latency following treatment (in seconds), “baseline” equals tail withdrawal latency prior to treatment (in seconds) and “cutoff” equals the investigator-imposed maximum trial time, which was 20 seconds in this experiment. Then, we applied simple linear regression to the ascending limb of the inverted-U-shaped dose-response curve (i.e., on data points up to, and including, the 0.1 mg/kg dose). Finally, we interpolated the dose at which %MPE = 50 (*i.e*., the half-maximal effect) from the linear regression curve. We applied an F-test to the ascending limb data at each time point to test for differences in slope or intercept of the linear regressions. To test for differences between BUP and BUP-D2 at each dose and overall, we conducted a two-way ANOVA with Sidak’s multiple comparisons test with data from all doses (*i.e*., ascending and descending limb) for each time point. In the pharmacokinetics experiment, blood concentrations of BUP, BUP-D2, and NorBUP versus time data from each rat were analyzed by non-compartmental analysis using WinNonlin Phoenix software (v. 8.3, Certara, Princeton, NJ). The pharmacokinetic calculations were dose-corrected and included half-life (t_1/2_), maximum concentration (C_max_), area under the curve (AUC), clearance, mean residence time (MRT), and steady state volume of distribution (V_ss_) for BUP and BUP-D2, and time to reach C_max_ (T_max_), C_max_, AUC, and MRT for NorBUP. Student’s two-tailed t-tests were applied using GraphPad Prism to determine whether these parameters differed for animals receiving BUP-D2 and those that received BUP. Welch’s t-test was performed in place of the Student’s t-test for NorBUP’s T_max_ and AUC because F-tests indicated that variances significantly differed between the BUP- and BUP-D2-treated groups.

## 3. RESULTS

### 3.1 Competition binding

Binding of BUP-D2 to opioid receptors with BUP-like affinity is crucial for BUP-D2 to retain the therapeutic effects of BUP. We used a competition receptor binding assay to determine and compare the affinities of BUP-D2 and BUP for human MOR, human DOR, and human KOR, and used morphine as a positive control. In these experiments, morphine and BUP exhibited affinity for each opioid receptor type similar to previously reported affinities^15,29^ (**Fig. 3**; **Table 1**). BUP-D2 and BUP exhibited sub-nanomolar affinity at each opioid receptor type (**Table 1**). BUP-D2 and BUP had significantly greater affinity for hMOR than did morphine, but BUP-D2 and BUP binding did not differ from each other at this opioid receptor (**Fig. 3A; Table 1**). Likewise, BUP-D2 and BUP affinities for hDOR did not differ from each other significantly, and both exhibited significantly greater affinity for hDOR than did morphine (**Fig. 3B; Table 1**). Finally, BUP-D2 and BUP affinities for hKOR did not differ from each other, and both had significantly greater affinity for hKOR than did morphine (**Fig. 3C; Table 1**).

**Figure 3.**
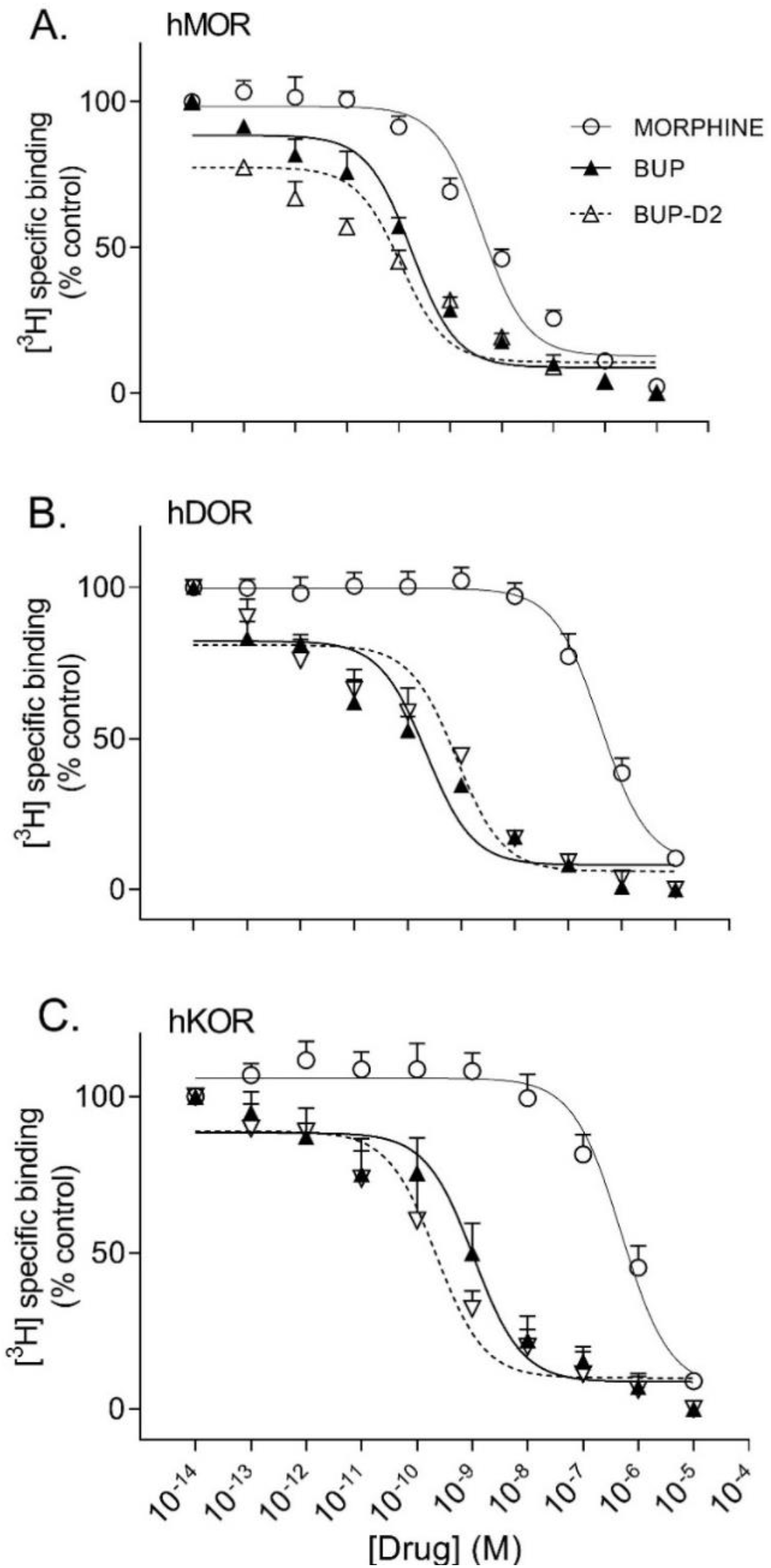
BUP-D2 and BUP have equal high affinity for opioid receptors. Data points and error bars represent mean and standard error of mean, respectively, of ^3^H-diprenorphine specific binding in homogenates containing hMOR (A), hDOR (B), or hKOR (C) as a percent of controls (y-axis) in the presence of varying concentrations (x-axis) of unlabeled morphine (circles), BUP (filled triangles), or BUP-D2 (unfilled triangles). “Control” refers to ^3^H-Diprenorphine binding in the presence of minimal unlabeled drug (10^-14^ M). n = 3-4 independent experiments performed in triplicate.

**Table 1.** Affinity of BUP-D2 for opioid receptors.

| Table 1. Affinity of BUP-D2 for opioid receptors |  |  |  |  |  |  |
| --- | --- | --- | --- | --- | --- | --- |
|  | Log K<sub>i</sub> values (M)<br>Mean (95% CI) |  |  | K<sub>i</sub> values (nM)<br>Mean (SEM) |  |  |
|  | hMOR | hDOR | hKOR | hMOR | hDOR | hKOR |
| **BUP-D2** | -10.3<br>(-11.1, -9.51) | -9.54<br>(-10.8, -8.26) | -10.2<br>(-11.7, -8.66) | 0.077<br>(0.038) | 0.424<br>(0.229) | 0.101<br>(0.494) |
| **BUP** | -10.0<br>(-10.2, -9.78) | -9.96<br>(-10.5, -9.45) | -9.55<br>(-11.1, -8.01) | 0.103<br>(0.016) | 0.119<br>(0.032) | 0.511<br>(0.342) |
| **Morphine** | **-8.66**<br>**(-9.07, -8.25)** | **-6.59**<br>**(-7.11, -6.07)** | **-6.71**<br>**(-7.60, -5.83)** | 2.50<br>(0.704) | 279<br>(83.5) | 239<br>(101) |
| K<sub>i</sub>, inhibitor constant; CI, confidence interval; SEM, standard error of mean; **Bolding** indicates significant difference from BUP and BUP-D2 (within receptor subtype), as determined by Tukey's multiple comparisons test (p < .005). Statistical analyses were conducted with log-transformed values only. n = 3-4 independent experiments conducted in duplicate. |  |  |  |  |  |  |

### 3.2 GTPγS Binding

After determining that BUP-D2 retains the binding affinity of BUP for opioid receptors, we wanted to determine whether BUP-D2 also retained the potency and intrinsic efficacy of BUP at opioid receptors. To accomplish this goal, we first measured [^35^S]GTPγS binding in homogenates containing each opioid receptor type in the presence or absence of a receptor-saturating concentration (1 μM) of BUP, BUP-D2, a positive control, or a negative control. The full agonists DAMGO, DPDPE, and U50,488 were used as positive controls for G-protein activation by hMOR, hDOR, and hKOR, respectively, and naltrexone (a non-selective opioid antagonist) was used as a negative control for all receptor types. Positive controls significantly activated their respective receptor by at least 175% of vehicle control levels (**Fig. 4**). As expected from previous reports^15,29^, BUP stimulated G-proteins in hMOR homogenates greater than baseline activity measured in vehicle controls, but less than positive control-stimulated activity (**Fig. 4A**), indicating that it acts as a partial agonist at hMOR. BUP-D2 likewise stimulated G-protein activation with partial agonist activity at hMOR (**Fig. 4A**). BUP and BUP-D2 did not stimulate G-proteins in hDOR homogenates and equally stimulated G-proteins in hKOR homogenates, suggesting that BUP and BUP-D2 both act as neutral antagonists of hDOR and partial agonists of hKOR (**Fig. 4B and 4C**). The negative control naltrexone did not stimulate G-proteins in hMOR or hDOR homogenates; however, it curiously stimulated G-proteins in hKOR homogenates with partial agonist activity.

**Figure 4.**
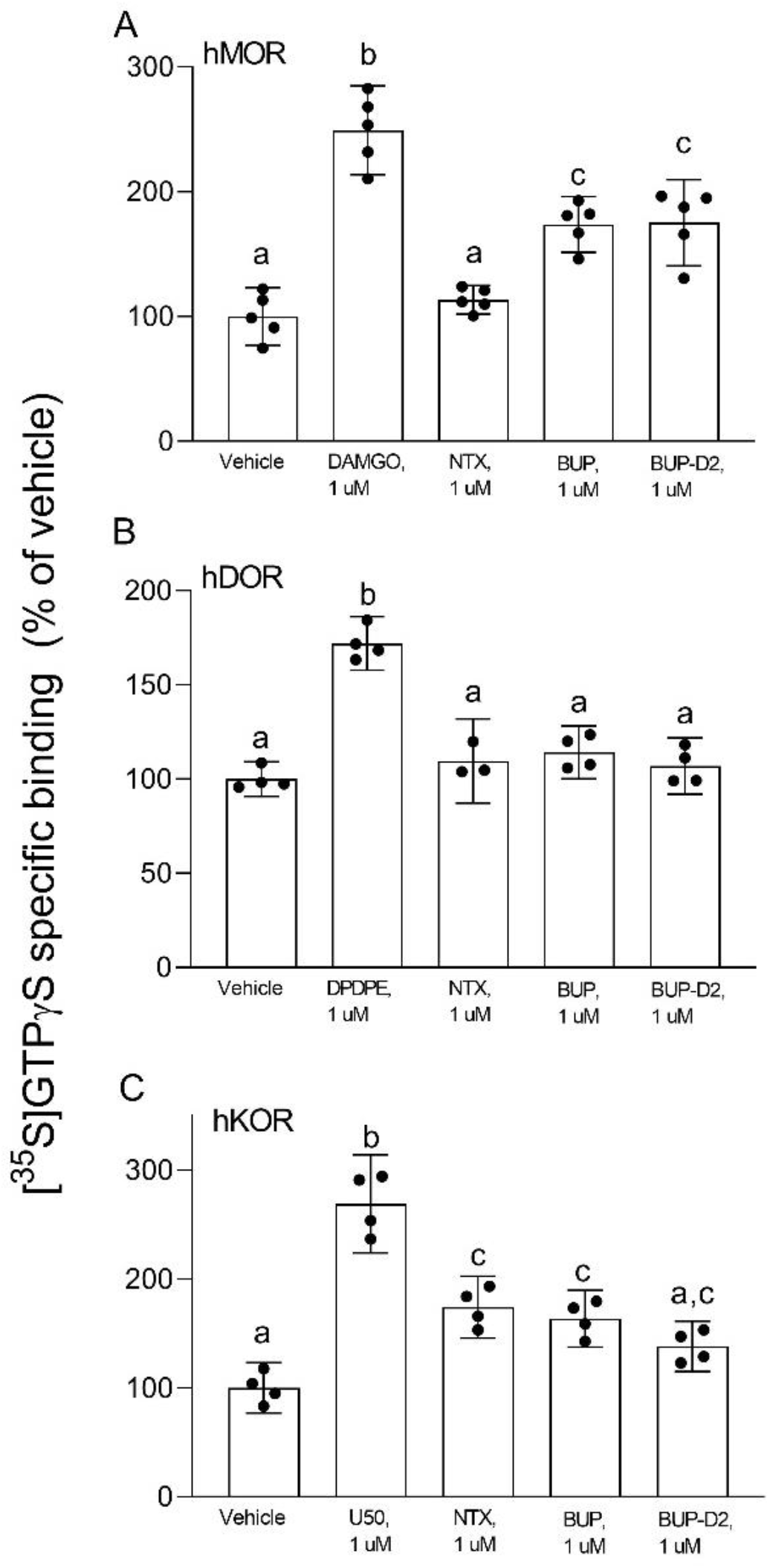
BUP-D2 and BUP activate opioid receptors with equal maximal efficacy. [^35^S]GTPγS specific binding (y-axis) represents G-protein activation in homogenates containing hMOR (A), hDOR (B), or hKOR (C) in the presence of vehicle (0.01% DMSO) or a receptor-saturating concentration (1 μM) of positive control full agonists, negative control antagonist naltrexone, BUP, or BUP-D2 (x-axis). Bars represent group means, closed circles represent values obtained from independent experiments, and error bars represent 95% confidence intervals. Within each panel, columns sharing letters are not significantly different (One-way ANOVA, Tukey’s multiple comparison test, p < 0.05, n = 3-5 independent experiments performed in triplicate).

To determine potency to activate opioid receptors, we next determined the concentration-effect of BUP and BUP-D_2_ to stimulate G-proteins in hMOR and hKOR homogenates (**Fig. 5**). Due to the lack of G-protein activation in screening experiments, concentration-effect curves of BUP and BUP-D2 were not generated using hDOR homogenates. BUP and BUP-D2 exhibited equal potency to activate G-proteins in hMOR homogenates and were both significantly more potent than the full agonist positive control DAMGO (**Fig. 5A; Table 2**). Likewise, in hKOR homogenates, BUP and BUP-D2 exhibited equal potency to activate G-proteins and were both significantly more potent than the full agonist positive control, U50,488 (**Fig. 5B; Table 2**). Greater potency of BUP to activate hMOR and hKOR relative to DAMGO and U50,488, respectively, was expected based on previous reports^15,29^. Maximal efficacies for BUP and BUP-D2 in the concentration-effect experiment were consistent with partial agonist efficacies determined in the screening experiment (**Fig. 4** vs. **Fig. 5**; **Table 2**), providing more evidence that BUP and BUP-D2 are partial agonists at hMOR and hKOR receptors.

**Figure 5.**
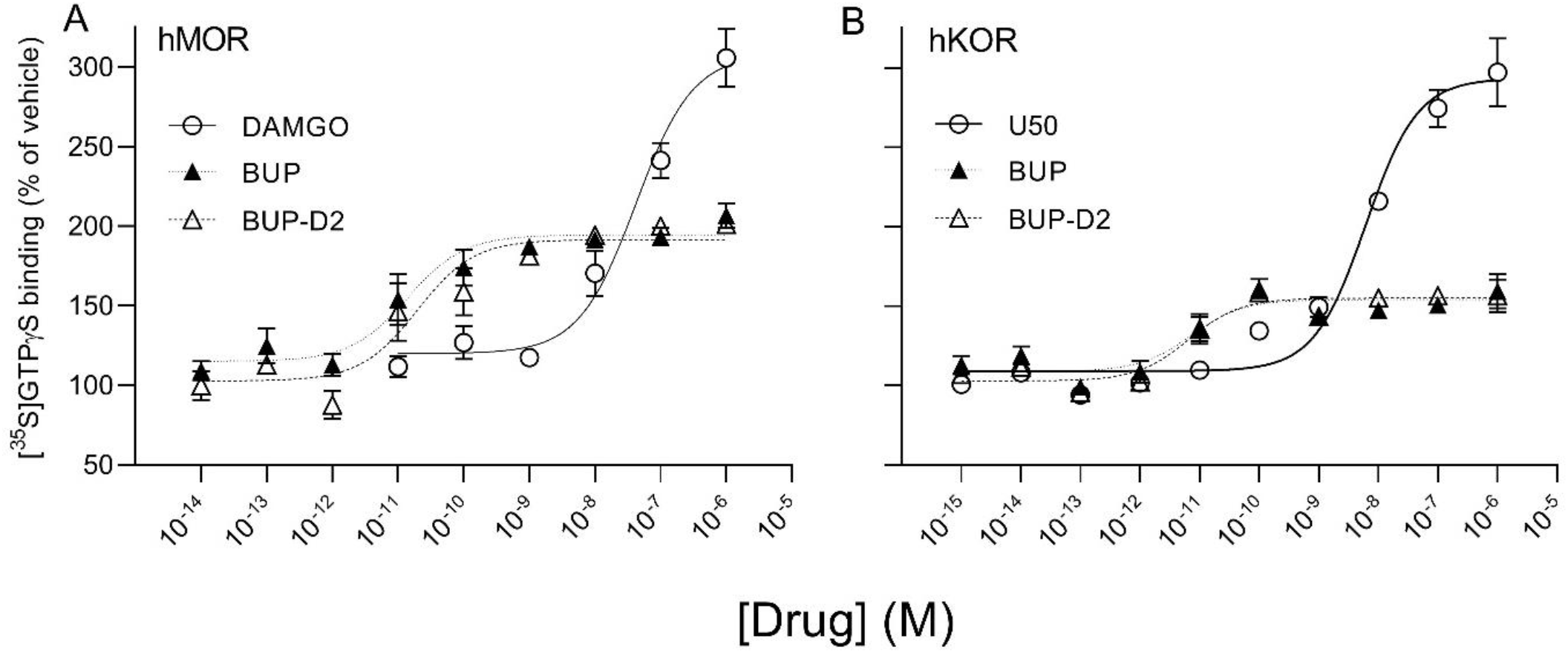
BUP-D2 and BUP activate hMOR and hKOR with equal, high potency. [^35^S]GTPγS specific binding (y-axis) represents G-protein activation in homogenates containing hMOR (A) or hKOR (B) in the presence of increasing concentrations (x-axis) of DAMGO (open circles, A), U50,488 (open circles, B), BUP (closed triangles), or BUP-D2 (open triangles). Symbols and error bars represent means and standard error of mean from independent experiments (n = 3-8 independent experiments per concentrations, conducted in triplicate).

**Table 2.**
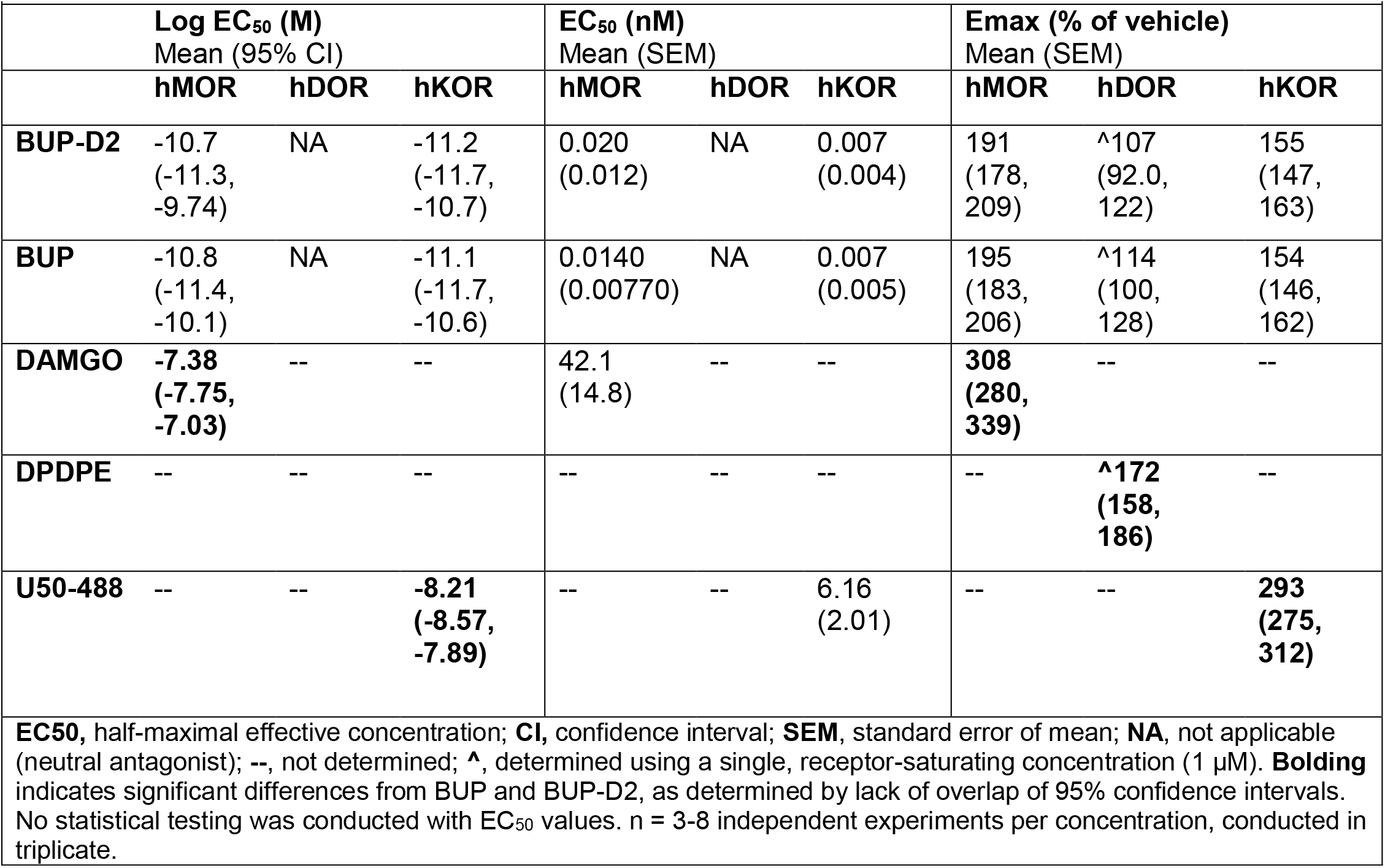
Potency and efficacy of BUP-D2 for opioid receptors.

### 3.3 Warm-water tail withdrawal analgesia assay

The antinociceptive effects of BUP and BUP-D2 were compared using the warm water tail withdrawal assay in female Long-Evans rats. In this experiment, we measured the latencies of rats to remove their tails from 50°C water 10 minutes and 60 minutes after an intravenous injection of either BUP or BUP-D2, up to a maximum cutoff of 20 seconds. Baseline latencies were equal for the BUP and BUP-D2 groups (**Fig. 6**). At both time points, there were no differences in tail withdrawal latency between BUP and BUP-D2 at any dose (**Fig. 6**). Likewise, there was no difference between BUP-D2 and BUP in ED50 values or percent maximum possible effect (**Table 3**). Maximal antinociception was achieved at 0.1 mg/kg at both time points for both drugs and declined at the higher dose of 0.3 mg/kg. This inverted U-shaped curve is atypical for most opioid agonists but is characteristic of BUP^30^, thus providing more evidence that BUP-D2 retains the opioid effects of BUP.

**Figure 6.**
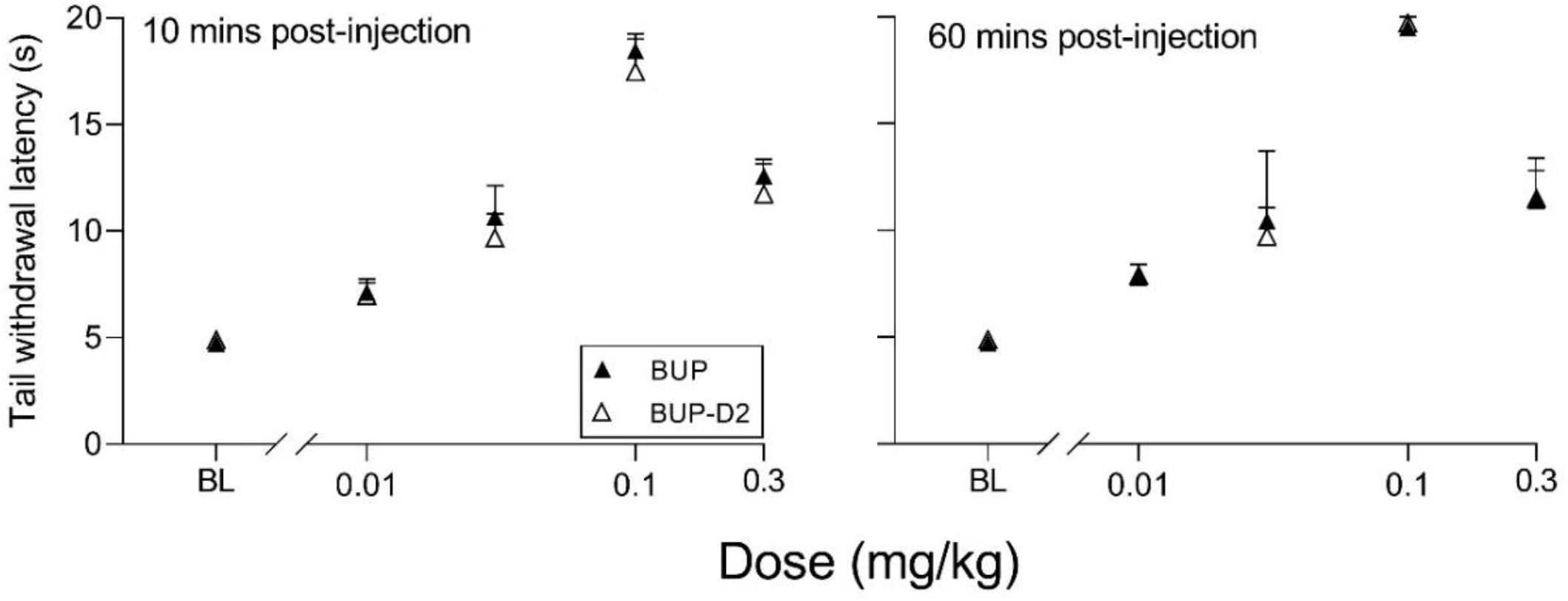
BUP-D2 retains the full antinociceptive potency and efficacy of BUP in the warm water tail withdrawal assay. Data points and error bars represent the mean and standard error, respectively, of the latency for rats to remove their tails from 50°C water (y-axis) before (baseline, “BL”) and after (1ü minutes, A, and 6ü minutes, B,) intravenous injection with BUP (closed triangles) or BUP-D2 (open triangles) at varying doses (x-axis). p > 0.05. two-way ANOVA with Sidak’s multiple comparisons test, n = 3-4 independent animals per dose.

**Table 3.** Potency and efficacy of BUP and BUP-D2 antinociception, mean (lower, upper 95% confidence interval)

|  | BUP | BUP-D2 |
| --- | --- | --- |
| **10 minutes** |  |  |
| ED<sub>50</sub> (mg/kg) | 0.052 (0.046, 0.058) | 0.056 (0.050, 0.065) |
| % MPE (at 0.1 mg/kg) | 90.1 (68.8, 111) | 84.4 (43.5, 125) |
| **60 minutes** |  |  |
| ED<sub>50</sub> (mg/kg) | 0.049 (0.040, 0.061) | 0.049 (0.045, 0.054) |
| % MPE (at 0.1 mg/kg) | 96.8 (83.2, 110) | 97.9 (89.1, 107) |

**ED50**, half-maximal effective dose; **% MPE**, percent maximum possible effect; slopes and intercepts were equal, as tested by F-test ( $P > 0.45$ ), and there was overlap of 95% confidence intervals between BUP and BUP-D2 for ED<sub>50</sub> and % MPE values at both time points. $n = 3-4$ independent animals per dose

### 3.4 Pharmacokinetics experiment

To determine how precision deuteration of BUP affects its pharmacokinetics, particularly blood concentrations of NorBUP *in vivo*, we measured rat blood concentrations of BUP, BUP-D2, and NorBUP following intravenous injection with either BUP or BUP-D2. Correcting for a difference in dose (0.9 mg/kg for BUP, 5.6 mg/kg for BUP-D2), we determined that mean C_max_ of NorBUP was over 19.6-fold lower for rats that received BUP-D2 relative to rats that received BUP (Figure 7A; Table 4). Although not statistically significant, AUC of NorBUP was over 10.8-fold lower, and T_max_ and MRT were over 2-fold higher, for BUP-D2-treated rats than BUP-treated rats (Table 4). Half-life, C_max_, AUC, and clearance of BUP-D2 and BUP did not differ (Table 5). However, MRT and steady state volume of distribution were increased for BUP-D2 relative to BUP (Table 5).

**Figure 7.**
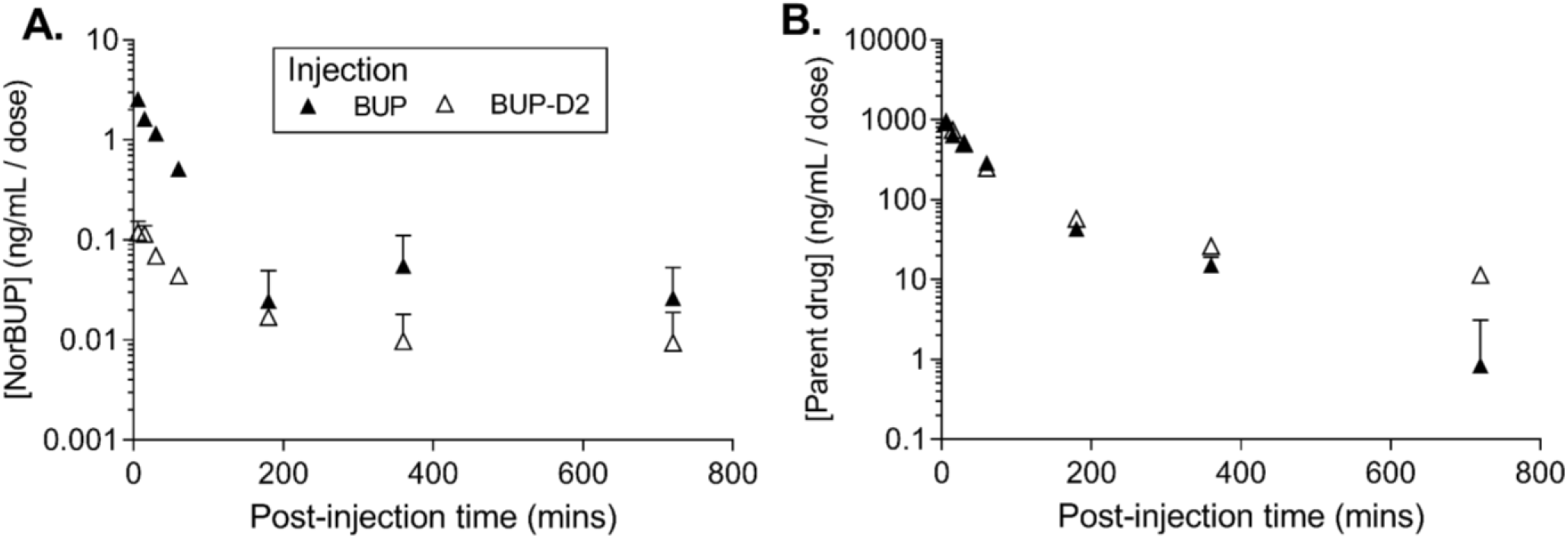
BUP-D2, relative to BUP, resists metabolism to NorBUP following bolus intravenous administration to rats. Dose-corrected blood concentration-time profiles of NorBUP (**A.**) and BUP or BUP-D2 (**B.**) following i.v. bolus injection of BUP (closed triangles) or BUP-D2 (open triangles). Symbols and error bars represent means and SEM from three rats.

**Table 4.** Dose-corrected pharmacokinetic parameters for NorBUP following administration of BUP and BUP-D2 mean (lower, upper 95% confidence interval)

|  | BUP | BUP-D2 | statistics |
| --- | --- | --- | --- |
| $T_{max}$, min | 6.00<br>(6.00, 6.00) | 12.0<br>(-0.908, 24.9) | $t(2) = 2.00$<br>$p = .183$ |
| $C_{max}$, ng/mL/mg/kg | **2.56**<br>**(2.26, 2.86)** | **0.130**<br>**(-0.003, 0.264)** | **$t(4) = 32.0$**<br>**$p < .0001$** |
| AUC, min*ng*kg/ mL*mg | 128<br>(-52.5, 309) | 11.9<br>(-0.445, 24.2) | $t(2.019) = 2.76$<br>$p = .109$ |
| MRT, min | 76.6<br>(-120, 273) | 156<br>(-130, 442) | $t(4) = 0.983$<br>$p = .381$ |

**Bolding** indicates statistically significant differences; $T_{max}$ , Time to reach maximum concentration; $C_{max}$ , maximum concentration; **AUC**, area under the curve; **MRT**, mean residence time

**Table 5.** Dose-corrected pharmacokinetic parameters for parent drugs BUP and BUP-D2 mean (lower, upper 95% confidence interval)

|  |  |  |  |
| --- | --- | --- | --- |
|  | **BUP** | **BUP-D2** | **statistics** |
| $t_{1/2}$, min | 101<br>(-56.9, 259.4) | 201<br>(18.0, 385) | $t(4) = 1.78$;<br>$p = .150$ |
| $C_{max}$, ng/mL/mg/kg | 1008<br>(111, 1905) | 790<br>(338, 1242) | $t(4) = 0.936$;<br>$p = .402$ |
| AUC, min*ng*kg/mL*mg | 60745.0<br>(13466.0, 108024) | 64485.0<br>(50762.0, 78207.0) | $t(4) = 0.327$;<br>$p = .760$ |
| CL, mL/min/kg | 17.6<br>(3.58, 31.7) | 15.6<br>(12.4, 18.8) | $t(4) = 0.610$<br>$p = .575$ |
| MRT, min | **74.5**<br>**(33.5, 115)** | **177**<br>**(56.2, 297)** | **$t(4) = 3.45$**<br>**$p = .026$** |
| $V_{ss}$, mL/kg | **1256**<br>**(825.6, 1687)** | **2731**<br>**(957.5, 4505)** | **$t(4) = 3.48$**<br>**$p = .025$** |
| **Bolding** indicates statistically significant differences; $t_{1/2}$, half-life; $C_{max}$, maximum concentration; **AUC**, area under the curve; **CL**, clearance; **MRT**, mean residence time; **Vss**, steady state volume of distribution | | | |

## 4. DISCUSSION

The present study is the first to apply the innovative strategy of precision deuteration to BUP and investigate this compound as a potentially novel therapeutic agent. The study demonstrates several important properties of BUP-D2: 1) that BUP-D2 retains BUP’s high affinity for opioid receptors; 2) that BUP-D2 retains BUP’s partial agonist efficacy for hMOR and hKOR and neutral antagonist efficacy for hDOR; 3) that BUP-D2 retains BUP’s high potency for hMOR and hKOR; 4) that BUP-D2 retains BUP’s antinociceptive potency and efficacy in rats; and 5) that NorBUP blood concentrations are lower in rats following intravenous administration of BUP-D2 relative to BUP. This is important evidence that BUP-D2 will retain the therapeutic activity of BUP and has the potential to reduce perinatal adverse effects (*e.g*., NOWS) by lowering maternal-fetal NorBUP exposure.

The unique therapeutic and safety profile of BUP, relative to methadone and other opioids, is thought to emerge from its high affinity for MOR, DOR, and KOR^15^, in combination with its high potency and partial agonist activity for MOR^15^. Determining the affinity of BUP-D2, relative to BUP for opioid receptors is the first step to determining whether BUP-D2 has potential as a therapeutic agent. Using a radioligand competition binding assay, we determined that BUP-D2 and BUP similarly displaced [^3^H]diprenorphine from each opioid receptor subtype, yielding equivalent sub-nanomolar affinities. Interestingly, [^3^H]diprenorphine binding in the presence of extremely low (femtomolar) concentrations of BUP and BUP-D2 was surprisingly lower than [^3^H]diprenorphine binding in the presence of vehicle controls, suggesting that BUP and BUP-D2 apparently displace [^3^H]diprenorphine at these concentrations. Additionally, unlike competition binding curves for the positive controls and morphine, the binding curves for BUP and BUP-D2 did not have a Hill slope that approximated −1. BUP and BUP-D2 instead displaced [^3^H]diprenorphine in a gradual linear manner throughout the entire range of concentrations used, with no clear minimum threshold for displacement. This behavior is possibly explained by differences in the dissociation rates of [^3^H]diprenorphine and BUP. Others have previously reported that after binding to opioid receptors, BUP dissociates from the receptor in ~40 minutes^31,32^. By comparison, diprenorphine and most other opioids dissociate from opioid receptors in 5-10 minutes^31–33^. The prolonged binding of BUP likely allows it to compete with [^3^H]diprenorphine at much lower concentrations than would be expected. Kinetic binding experiments specifically designed to measure dissociation rates are needed to definitively confirm that BUP-D2 retains the slow dissociation rate of BUP. In any case, BUP-D2 exhibiting atypical receptor binding similar to that of BUP at low concentrations is further evidence that precision deuteration does not alter the opioid receptor binding properties of BUP.

After confirming that receptor affinity for BUP-D2 was unchanged, relative to BUP, we next determined the potency and efficacy of BUP-D2 relative to BUP. It is important for BUP-D2 to retain the partial agonist efficacy of BUP, such that it stimulates MORs enough to prevent withdrawal (following proper induction) without the high risk of diversion, abuse, respiratory depression, and death observed with higher efficacy opioids. We used both *in vitro* and *in vivo* models to compare the potency and efficacy of BUP-D2 to BUP. Our *in vitro* approach measured BUP- and BUP-D2-induced G-protein stimulation in homogenates of CHO cells that were stably transfected with human MOR, DOR, and KOR, and determined that BUP-D2 and BUP stimulated G-proteins with equal potency and efficacy. More studies are needed to evaluate the potency and efficacy of BUP-D2 for other pathways downstream of G-protein activation that may affect its pharmacology at the whole-organism level, including β-arrestin-2 recruitment, and modulation of mitogen-activated protein kinase activity and gating of G-protein coupled inward rectifying potassium channels, as well as adenylyl cyclase activity and cAMP-modulated calcium channel opening and intracellular calcium concentrations. However, our results of the warm water tail withdrawal assay in rats provides the first evidence that BUP-D2 retains potency and efficacy similar to that of BUP for antinociception at the *in vivo* level. Importantly, BUP-D2 exhibited an inverted U-shaped curve that is distinctive of BUP^30^. Increasing antinociception at doses on the ascending limb of the dose-response curve predominantly results from MOR activation^34^. However, at higher doses, activation of nociception/orphanin FQ receptors by BUP is thought to antagonize MOR-induced antinociception, causing the descending limb of the dose-response curve^35^. The close matching of both ascending and descending limbs for BUP and BUP-D2 is compelling evidence that precision deuteration of BUP does not alter these pharmacodynamic effects.

The pharmacokinetics of BUP-D2 and BUP differed. The MRT, which is the average time a molecule of drug spends in the body, was over 1.5 hours longer for BUP-D2 than for BUP, suggesting that BUP-D2 resists metabolism and/or clearance relative to BUP. However, there was no difference in half-life, C_max_, AUC, or clearance between BUP-D2 and BUP. The differences in MRT are probably not due to different doses of BUP-D2 and BUP being administered because previous rat studies demonstrated that the MRT of BUP is dose-independent^36,37^, even over a wide range of 1 to 30 mg/kg^36^. Our study also determined that V_ss_, which describes the propensity of a drug to leave the blood and distribute to other tissues at steady state, was higher for BUP-D2 than for BUP, indicating that BUP-D2 was more readily distributed outside of blood than BUP. This effect may be partially or completely explained by the differences in dosing of BUP-D2 and BUP, because V_ss_ of BUP was previously determined to increase with increasing dose^36^. Investigators of that study attributed the dose-dependent increase in V_ss_, as well as an increase in clearance, to increased NorBUP concentrations competing against BUP for plasma protein binding sites, leading to higher concentrations of free BUP that could distribute from plasma. Our study does not support this explanation because we observed lower NorBUP concentrations following BUP-D2 administration, even without application of a dose correction. The contributions of metabolic pathways other than CYP-mediated *N*-dealkylation, such as glucuronidation, should be considered for their potential to compensate for resistance to the NorBUP pathway. As such, we expect that blood concentrations of glucuronide metabolites of BUP-D2 will be elevated relative to blood concentrations of glucuronide metabolites of BUP. Our ongoing work examines the effects of BUP deuteration on its glucuronidation.

The lower blood concentrations of NorBUP following intravenous administration of BUP-D2 relative to BUP further suggests that these drugs are metabolized differently. It is unlikely that dosing differences explain the lower NorBUP blood concentrations (*e.g*., by greater auto-inhibition of BUP-D2 metabolism due to the higher dose of BUP-D2 compared to BUP) because NorBUP blood concentrations increase in a linear manner with increasing BUP dosing in rats^36^. Although this work provides proof of concept that BUP-D2 resists metabolism to NorBUP, more work is needed to determine the clinical relevance of the NorBUP reduction and whether treatment with BUP-D2 during pregnancy would lower risk and severity of NOWS relative to treatment with BUP. Studies examining pharmacokinetics and NOWS liability of BUP-D2 during chronic administration and pregnancy would help elucidate the clinical relevance of BUP. These studies would determine whether such a reduction in maternal plasma concentrations can lower total fetal opioid exposure enough to reduce NOWS. Additionally, although NorBUP is thought to contribute little to therapeutic effects of treatment due to its low brain penetrance, studies are needed to empirically confirm that reduction of NorBUP does not have a negative impact on the effectiveness of maternal treatment.

In conclusion, this innovative study takes the important first steps for developing a potentially improved treatment for OUD during pregnancy. We envision that BUP-D2 will effectively treat maternal OUD similar to BUP, but will improve upon BUP by lowering the total cumulative opioid exposure endured by the fetus throughout gestation. Because maternal plasma concentrations of a drug are a major determinant of fetal exposure to the drug^38,39^, reducing maternal plasma concentrations of NorBUP by interfering with NorBUP formation is likely to reduce fetal exposure to NorBUP and thereby mitigate negative effects of maternal treatment of OUD.

## Abbreviations

AUC: area under the curve
BUP: buprenorphine
BUP-D2: deuterated buprenorphine
CHO: Chinese Hamster Ovary cells
CL: clearance
Cmax: maximum serum concentration
DAMGO: [D-Ala^2^, *N-* MePhe^4^, Gly-ol]-enkephalin
DPDPE: [D-Pen^2^,D-Pen^5^]enkephalin
DPMs: decays per minute
GDP: guanosine diphosphate
GTPγS: guanosine 5’-O-[gamma-thio]triphosphate
hDOR: human delta opioid receptors
hMOR: human mu opioid receptors
hKOR: human kappa opioid receptors
LC/MS/MS: liquid chromatography with tandem mass spectrometry
MPE: maximum possible effect
MRT: mean residence time
NorBUP: norbuprenorphine
NOWS: neonatal opioid withdrawal syndrome
t_1/2_: half-life
TLC: thin layer chromatography
T_max_: time to maximum serum concentration
OUD: opioid use disorder

## Acknowledgements and role of the funding sources

We acknowledge and thank the National Institute on Drug Abuse Drug Supply Program for providing norbuprenorphine for this study. Research reported in this publication was supported by the National Institutes of Health (NIH) National Center for Advancing Translational Sciences (NCATS) under award numbers UL1 TR003107, U54TR001629 and KL2TR000063 and by the NIH National Institute on Drug Abuse (NIDA) under award numbers R21DA049585 and T32DA022981. Additional support was provided by the American Society for Pharmacology and Experimental Therapeutics through a Summer Undergraduate Research Fellowship. The funding sources had no involvement in the preparation of the manuscript or the decision to submit it for peer review and publication. The content is solely the responsibility of the authors and does not necessarily represent the official views of the National Institutes of Health.

## Disclosures

Dr. Moran is the CEO and has equity interest in PinPoint Testing, LLC, a UAMS/Bioventures spinoff company that is accredited for clinical and forensic testing of whole blood. Dr. Alund was an employee of PinPoint Testing, LLC, at the time the experiments were conducted. Drs. Janganati, Crooks, and Brents are inventors on a provisional patent entitled “BUP-D2 as a Protective Agent for Fetal Subjects Against Full-Agonist Opioid Exposure.” All other authors declare that the research was conducted in the absence of any commercial or financial relationships that could be construed as a potential conflict of interest.

## Author contributions

**Janganati**: conceptualization, methodology, validation, formal analysis, investigation, writing—review and editing; **Salazar**: formal analysis, investigation, writing—original draft; **Parks**: investigation, writing—review and editing; **Gorman:** investigation, writing—review and editing; **Prather:** conceptualization, resources, writing—review and editing, funding acquisition; **Peterson**: formal analysis, writing—review and editing; **Alund:** methodology, formal analysis, investigation, writing—review and editing; **Moran:** methodology, formal analysis, resources, writing—review and editing, supervision, funding acquisition; **Crooks**: conceptualization, methodology, resources, writing—review and editing, supervision, funding acquisition; **Brents:** conceptualization, formal analysis, investigation, writing—review and editing, supervision, project administration, funding acquisition.

## Notes

### Summary of Updates

Addition of rat pharmacokinetics data demonstrating that plasma concentrations of norbuprenorphine are lower following intravenous injection with deuterated buprenorphine relative to buprenorphine.

## REFERENCES

1. Hirai, A. H.; Ko, J. Y.; Owens, P. L.; Stocks, C.; Patrick, S. W. JAMA 2021, 325, 146.

2. Haight, S. C.; Ko, J. Y.; Tong, V. T.; Bohm, M. K.; Callaghan, W. M. MMWR Morb. Mortal. Wkly. Rep. 2018, 67, 845.

3. Jansson, L. M.; Velez, M.; Harrow, C. J. Opioid Manag. 2009, 5, 47.

4. Strahan, A. E.; Guy, G. P.; Bohm, M.; Frey, M.; Ko, J. Y. JAMA pediatrics 2020, 174, 200.

5. Maeda, A.; Bateman, B. T.; Clancy, C. R.; Creanga, A. A.; Leffert, L. R. Anesthesiology 2014, 121, 1158.

6. Brogly, S. B.; Velez, M. P.; Werler, M. M.; Li, W.; Camden, A.; Guttmann, A. Epidemiology 2021, 32, 448.

7. Azuine, R. E.; Ji, Y.; Chang, H.; Kim, Y.; Ji, H.; DiBari, J.; Hong, X.; Wang, G.; Singh, G. K.; Pearson, C. JAMA network open 2019, 2, e196405.

8. Nørgaard, M.; Nielsson, M. S.; Heide-Jørgensen, U. Substance abuse: research and treatment 2015, 9, SART. S23547.

9. Krans, E. E.; Kim, J. Y.; Chen, Q.; Rothenberger, S. D.; James, A. E.; Kelley, D.; Jarlenski, M. P. Addiction 2021, 116, 3504.

10. Jones, H. E.; Kaltenbach, K.; Heil, S. H.; Stine, S. M.; Coyle, M. G.; Arria, A. M.; O’Grady, K. E.; Selby, P.; Martin, P. R.; Fischer, G. N. Engl. J. Med. 2010, 363, 2320.

11. Bartu, A. E.; Ilett, K. F.; Hackett, L. P.; Doherty, D. A.; Hamilton, D. Aust. N. Z. J. Obstet. Gynaecol. 2012, 52, 342.

12. Deshmukh, S. V.; Nanovskaya, T. N.; Ahmed, M. S. J. Pharmacol. Exp. Ther. 2003, 306, 1099.

13. Chang, Y.; Moody, D. E.; McCance-Katz, E. F. Drug Metab. Dispos. 2006, 34, 440.

14. Kobayashi, K.; Yamamoto, T.; Chiba, K.; Tani, M.; Shimada, N.; Ishizaki, T.; Kuroiwa, Y. Drug Metab. Dispos. 1998, 26, 818.

15. Huang, P.; Kehner, G. B.; Cowan, A.; Liu-Chen, L. Y. J. Pharmacol. Exp. Ther. 2001, 297, 688.

16. Shah, D.; Brown, S.; Hagemeier, N.; Zheng, S.; Kyle, A.; Pryor, J.; Dankhara, N.; Singh, P. Springerplus 2016, 5, 854.

17. Kacinko, S. L.; Jones, H. E.; Johnson, R. E.; Choo, R. E.; Huestis, M. A. Clin. Pharmacol. Ther. 2008, 84, 604.

18. Concheiro, M.; Jones, H. E.; Johnson, R. E.; Choo, R.; Shakleya, D. M.; Huestis, M. A. Ther. Drug Monit. 2010, 32, 206.

19. Jones, H. E.; Dengler, E.; Garrison, A.; O’Grady, K. E.; Seashore, C.; Horton, E.; Andringa, K.; Jansson, L. M.; Thorp, J. Drug Alcohol Depend. 2014, 134, 414.

20. O’Connor, A. B.; O’Brien, L.; Alto, W. A. Eur. Addict. Res. 2016, 22, 127.

21. Griffin, B. A.; Caperton, C. O.; Russell, L. N.; Cabalong, C. V.; Wilson, C. D.; Urquhart, K. R.; Martins, B. S.; Zita, M. D.; Patton, A. L.; Alund, A. W.; Owens, S. M.; Fantegrossi, W. E.; Moran, J. H.; Brents, L. K. J. Pharmacol. Exp. Ther. 2019, 1, 9–17.

22. Brown, S. M.; Campbell, S. D.; Crafford, A.; Regina, K. J.; Holtzman, M. J.; Kharasch, E. D. J. Pharmacol. Exp. Ther. 2012, 343, 53.

23. Alhaddad, H.; Cisternino, S.; Decleves, X.; Tournier, N.; Schlatter, J.; Chiadmi, F.; Risede, P.; Smirnova, M.; Besengez, C.; Scherrmann, J. M.; Baud, F. J.; Megarbane, B. Crit. Care Med. 2012, 40, 3215.

24. Miwa, G. T.; Walsh, J. S.; Kedderis, G. L.; Hollenberg, P. F. J. Biol. Chem. 1983, 258, 14445.

25. Harbeson, S. L.; Morgan, A. J.; Liu, J. F.; Aslanian, A. M.; Nguyen, S.; Bridson, G. W.; Brummel, C. L.; Wu, L.; Tung, R. D.; Pilja, L.; Braman, V.; Uttamsingh, V. J. Pharmacol. Exp. Ther. 2017, 362, 359.

26. Uttamsingh, V.; Gallegos, R.; Liu, J. F.; Harbeson, S. L.; Bridson, G. W.; Cheng, C.; Wells, D. S.; Graham, P. B.; Zelle, R.; Tung, R. J. Pharmacol. Exp. Ther. 2015, 354, 43.

27. Yadlapalli, J. S. K.; Ford, B. M.; Ketkar, A.; Wan, A.; Penthala, N. R.; Eoff, R. L.; Prather, P. L.; Dobretsov, M.; Crooks, P. A. Pharmacological research 2016, 113, 335.

28. Cheng, Y.; Prusoff, W. H. Biochem. Pharmacol. 1973, 22, 3099.

29. Olson, K. M.; Duron, D. I.; Womer, D.; Fell, R.; Streicher, J. M. PloS one 2019, 14, e0217371.

30. Christoph, T.; Kogel, B.; Schiene, K.; Meen, M.; De Vry, J.; Friderichs, E. Eur. J. Pharmacol. 2005, 507, 87.

31. Olofsen, E.; Algera, M. H.; Moss, L.; Dobbins, R. L.; Groeneveld, G. J.; van Velzen, M.; Niesters, M.; Dahan, A.; Laffont, C. M. JCI insight 2022, 7(9), e156973.

32. Sadee, W.; Rosenbaum, J.; Herz, A. J. Pharmacol. Exp. Ther. 1982, 223, 157.

33. Cunningham, V. J.; Hume, S. P.; Price, G. R.; Ahier, R. G.; Cremer, J. E.; Jones, A. K. Journal of Cerebral Blood Flow & Metabolism 1991, 11, 1.

34. Yamamoto, T.; Shono, K.; Tanabe, S. J. Pharmacol. Exp. Ther. 2006, 318, 206.

35. Lutfy, K.; Eitan, S.; Bryant, C. D.; Yang, Y. C.; Saliminejad, N.; Walwyn, W.; Kieffer, B. L.; Takeshima, H.; Carroll, F. I.; Maidment, N. T. Journal of Neuroscience 2003, 23, 10331.

36. Gopal, S.; Tzeng, T.; Cowan, A. European journal of pharmaceutical sciences 2002, 15, 287.

37. Ohtani, M.; Kotaki, H.; Uchino, K.; Sawada, Y.; Iga, T. Drug Metab. Dispos. 1994, 22, 2.

38. Anoshchenko, O.; Storelli, F.; Unadkat, J. D. Drug Metab. Dispos. 2021, 49, 919.

39. Zhang, Z.; Unadkat, J. D. Drug Metab. Dispos. 2017, 45, 939.

