## Supporting data for "Deuterated buprenorphine retains pharmacodynamic properties of buprenorphine and resists metabolism to the active metabolite norbuprenorphine in rats"

Figure 1.

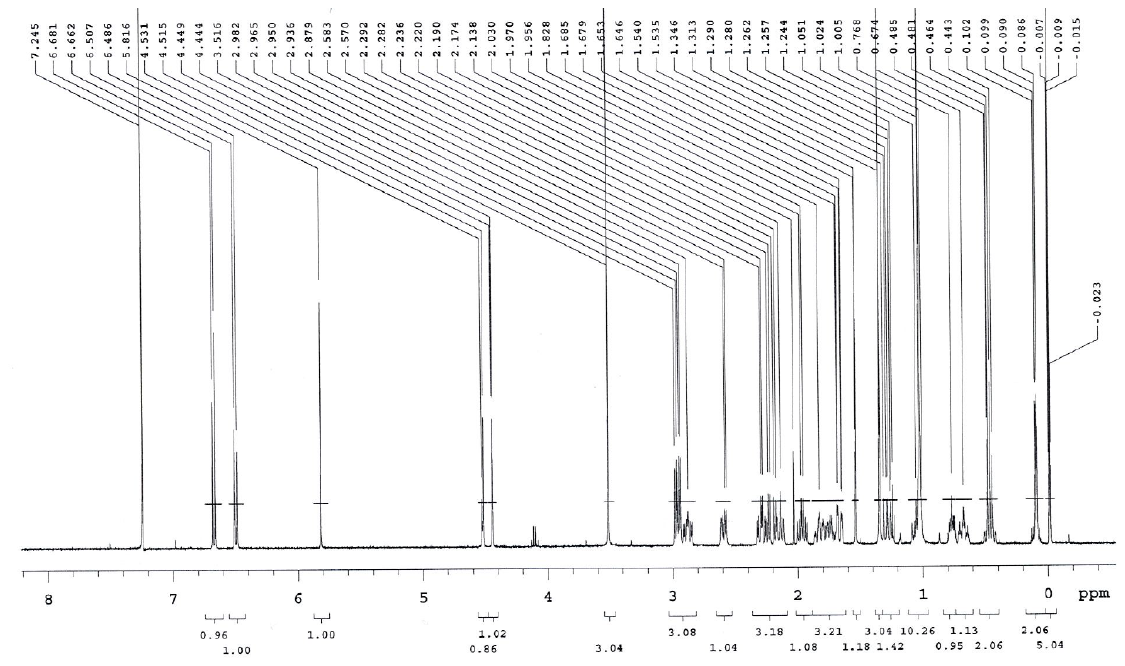
A.

1H-NMR spectroscopy of BUP-D2

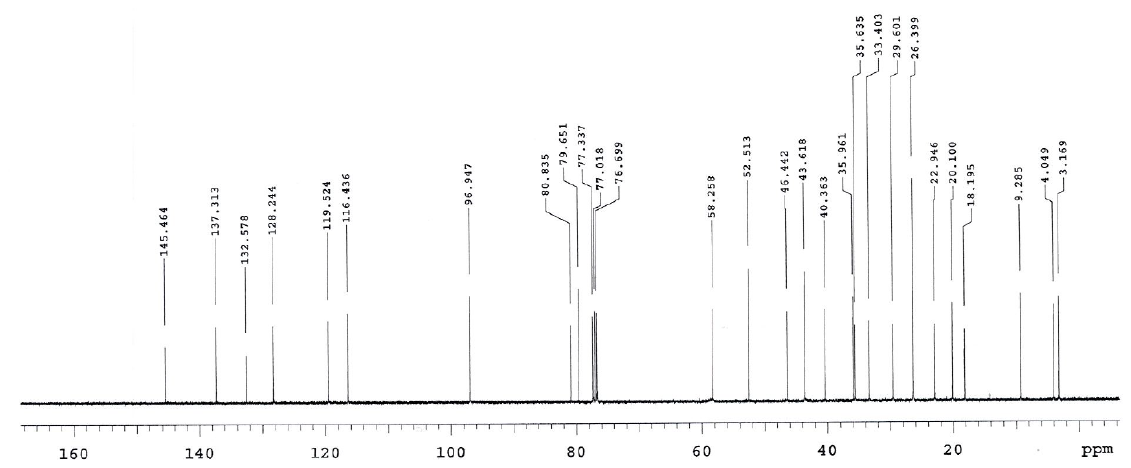
B.

13C-NMR spectroscopy of BUP-D2

Figure 2.

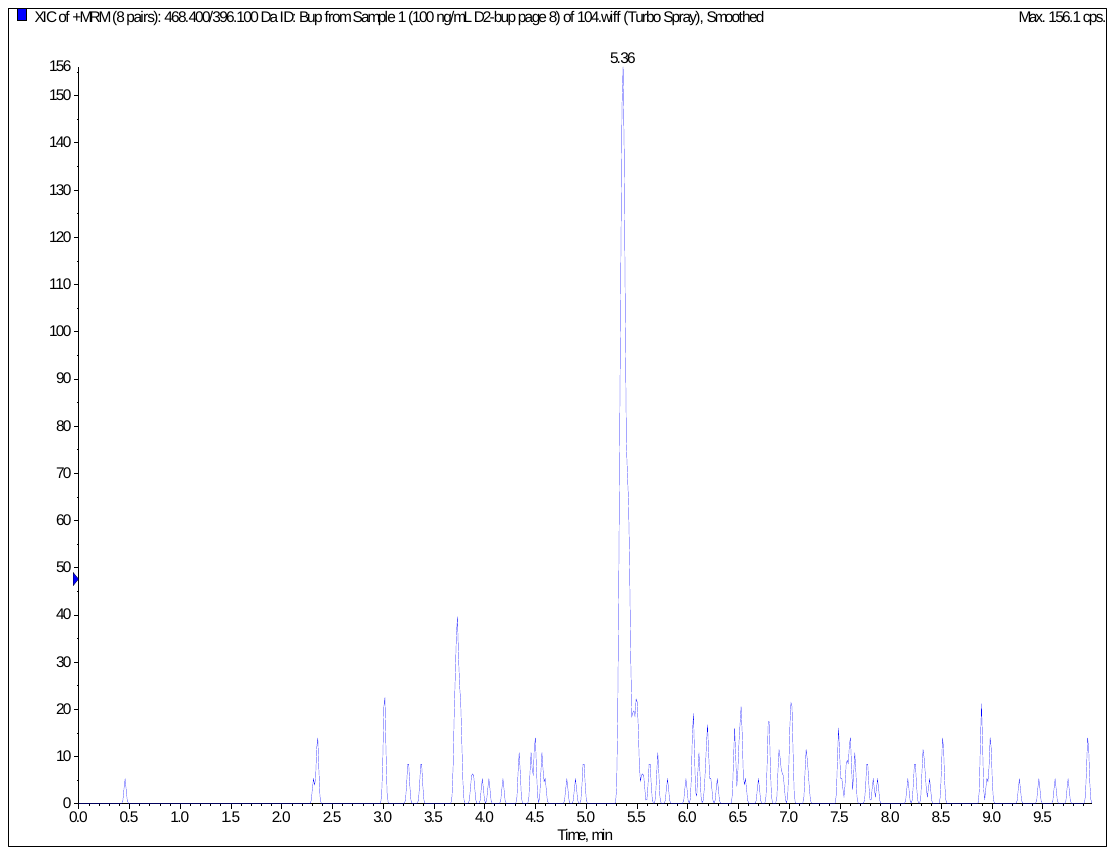

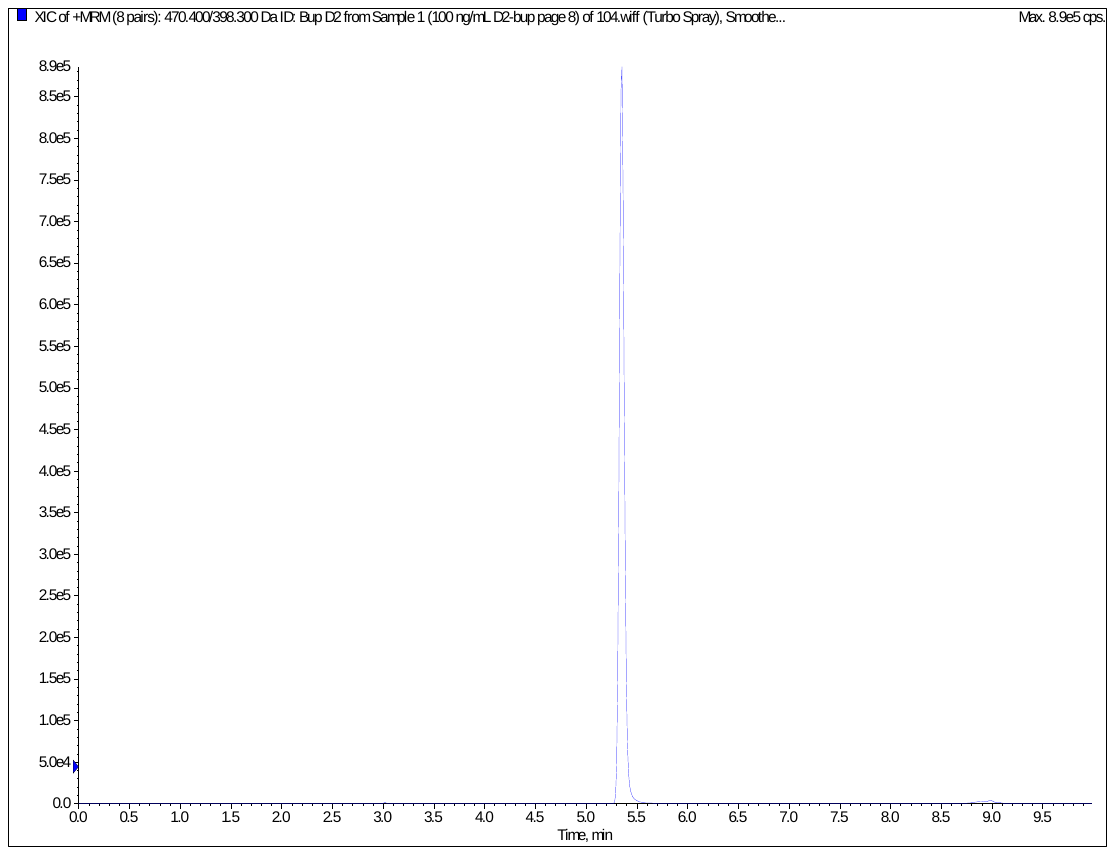

MRM chromatogram of 100 ng/mL D2 buprenorphine (top is transition for Bup m/z 468.4🡪396.1 and bottom is transition for BupD2 m/z 470.4🡪398.3).
